# *Anopheles* mosquitoes exposed to long-acting antimalarials via drug-spiked bloodmeal absorb drug but do not suffer fitness costs

**DOI:** 10.1101/2025.11.13.688204

**Authors:** Morgan M. Goheen, Alessandra Orfanó, S. Rodrigue Dah, Brian D. Foy, Doug E. Brackney, Fangyong Li, Jean-Bosco Ouédraogo, Dari F. Da, Roch K. Dabiré, A. Fabrice Somé, Amy K. Bei, Sunil Parikh

**Affiliations:** Section of Infectious Diseases, Yale School of Medicine, New Haven, Connecticut, United States of America; Department of Epidemiology of Microbial Diseases, Yale School of Public Health, New Haven, Connecticut, United States of America; Institut de Recherche en Sciences de la Santé, Direction Regionale de l’Ouest, Bobo-Dioulasso, Burkina Faso; Department of Microbiology, Immunology, and Pathology, College of Veterinary Medicine and Biomedical Sciences, Colorado State University, Fort Collins, Colorado, United States of America; Connecticut, Agricultural Experiment Station, New Haven, Connecticut, United States of America; Yale Center for Analytical Sciences, Yale School of Public Health, New Haven, Connecticut, United States of America; Institut des Sciences et Techniques, Bobo-Dioulasso, Burkina Faso

## Abstract

The World Health Organization’s recommendations regarding the use of antimalarials for the prevention of malaria in endemic areas have greatly expanded, allowing more flexibility in the demographic groups and regions where chemoprevention and mass treatment are acceptable. An overlooked aspect of expanding human population-level drug exposure is the downstream impact of ingested drug on the mosquito vector. Data suggest both infected and uninfected *Anopheles* mosquitoes re-feed often with ≥4 blood meals during their lifespan. This provides repeated opportunities for mosquitoes to ingest drug via bloodmeals taken from people with antimalarials in the bloodstream and raises questions as to whether exposure may impact the mosquito itself, and/or parasites developing within the infected mosquito. We investigated the impact of exposure to physiologic levels of commonly used long-acting human antimalarials in the *Anopheles* mosquito via drug-spiked blood feeds. We did not observe any significant differences in mosquito feeding, behavior, fertility, or viability after ingestion of amodiaquine, desethylamodiaquine, piperaquine, and sulfadoxine-pyrimethamine in either lab-reared *An. gambiae* or field-derived *An. coluzzii* mosquitoes. Interrogating drug distribution within mosquitoes utilizing LC-MS/MS, desethylamodiaquine, the longer-acting active metabolite of amodiaquine, was fed at 2 concentrations (1/2X and 2X C_max_) with drug subsequently detected in a dose-dependent manner in pooled whole mosquitoes, midguts and hemolymph. This was significant for whole mosquitoes harvested at 24hrs and 120hrs, and midguts harvested at 24hrs. Between 24 to 120 hrs, drug decreased in midguts but increased in hemolymph. Our results show biochemical evidence of antimalarial absorption into *Anopheles* hemolymph following bloodmeal ingestion. These studies lay the foundation for future work to assess the impact of vector-stage antimalarial drug exposure on parasite progression throughout development in the mosquito, which could in turn have important implications for transmission dynamics and drug resistance spread.

**Author summary:** Drug resistance to first-line antimalarials has emerged in multiple African countries. A better understanding of antimalarial drug resistance emergence and spread is critical in preventing further morbidity and mortality. Millions regularly receive antimalarials for prophylaxis and mass treatment that are purposefully long-acting. *Anopheles* mosquitoes re-feed frequently, and these malaria vectors (both infected and uninfected) routinely feed on people whose blood contains these long-acting antimalarials. Parasites take approximately 10 days to develop within the mosquito. There is published precedent that several antimalarials can act upon vector-stage parasites, yet any potential impact of antimalarials on mosquitoes and/or parasites developing within has been largely overlooked. We questioned whether antimalarials ingested in mosquito bloodmeals could influence parasite development and drug resistance selection. As initial investigations, we exposed uninfected *Anopheles* mosquitoes to commonly used long-acting antimalarials via drug-spiked bloodmeals, mimicking predicted physiologic drug exposure. Investigated drugs did not impact mosquito viability. However, mass spectrometry confirmed drug absorption in whole mosquitoes, as well as within midguts and circulatory fluid, several days after feeding, demonstrating that mosquitoes can ingest key drugs without suffering fitness costs, and these drugs can persist in mosquitoes. This highlights the potential for antimalarials to impact parasite development and drug resistance selection within the mosquito.

## Introduction

Despite gains in detection, treatment, and prevention strategies over previous decades, including recent approval of the first malaria vaccines, malaria remains a human pathogen of significant consequence, causing 597,000 deaths and 263 million cases in 2023 [1]. Unfortunately, formerly decreasing morbidity and mortality trends have now flatlined – particularly worrisome as insecticide resistance in the *Anopheles* mosquito vector and drug resistance in the predominant *Plasmodium falciparum* parasite species rapidly emerges and spreads [2–5]. Malaria control efforts are necessarily multipronged and include the widespread use of long-acting antimalarials for both chemoprevention and treatment. In the West African Sahel region, seasonal malaria chemoprevention (SMC) has been deployed to children under 5 years old as prevention during high transmission periods since ∼2014. Thus far, administration of the long-acting drugs sulfadoxine (SDX)-pyrimethamine (PYR) (referred to as SP together) plus amodiaquine (AQ) (SP-AQ combined for SMC) has been the sole SMC regimen, and was administered to 53 million children per monthly cycle in 2023 [1,6–8] in countries with seasonal malaria transmission. SP-AQ for SMC is now also being considered for use in older children. Other WHO-endorsed long-acting prophylaxis and disease reduction strategies include intermittent preventative treatment in pregnancy (IPTp; all pregnant women in malaria endemic areas), which also employs SP (though dihydroartemisinin (DHA) + piperaquine (PPQ) (DP in combination) is also being considered), and mass drug administration (MDA; community wide treatment in moderate-high transmission areas to reduce disease burden), which typically involves DP. First-line antimalarial treatment typically combines a short-acting artemisinin derivative with a long-acting partner (e.g. artesunate-AQ (ASAQ), artemether-lumefantrine (AL), or DP [6].

Recommendations for community antimalarial dosing in the context of prophylaxis and mass treatment are now expanding, which is a “paradigm shift” according to the WHO because its guidelines no longer specify strict age groups, transmission intensity thresholds, numbers of doses or cycles, or specific drugs [6]. Given the intentional long-acting nature of many antimalarials, with expanded use, it is anticipated that large proportions of a population could have drugs in their circulatory system for a majority of the transmission season(s). In fact, there is already abundant evidence of widespread circulating levels of human antimalarials in malaria endemic areas. Our group demonstrated that 90% of Ugandan children treated with DP had PPQ detectable in plasma at their next malaria episode, even those occurring up to 126 days post-treatment [9]. A Tanzanian study found that 74% of people had ≥1 antimalarial detectable despite no reported intake for 28 days prior [10]. The impact of such high levels of extended drug presence in human populations raises inevitable questions regarding how such exposure could facilitate drug resistance emergence and/or spread.

Malaria drug resistance is generally thought to arise through mutation and selection during symptomatic human blood stage infection, when both parasite replication and drug pressure are greatest [11]. In contrast, mass drug intervention is thought to reduce symptomatic cases and transmission, and in turn, pose a lower risk of resistance emergence and selection [12,13]. However, very little is known about how drugs ingested by mosquitoes during bloodmeals could impact susceptible parasite forms in the mosquitoes, as well as parasite resistance selection within the vector [14].

Revisiting the nuances of vector feeding habits and parasite development within the vector (known as sporogony) is central to this question. *Anopheles spp.* mosquitoes become infected after feeding on a person harboring gametocytes (the *Plasmodium* sexual stage); in the vector midgut parasites undergo sexual replication, with subsequent oocysts beginning to form within ∼24 hours. Oocysts then further develop under the mosquito midgut epithelium for ∼9-14 days before releasing thousands of sporozoites which migrate via hemolymph (circulatory fluid) to salivary glands for entry into a new host upon next feed [15–17]. However, mosquitoes do not only feed twice during their lifetime [18–21]. The *Anopheles* vector can survive up to ∼20 days and have multiple egg laying cycles in the wild. They re-feed during each gonadotrophic cycle and sometimes within a single gonadotrophic cycle (particularly given nutrient demands and feeding disruptions) [18,21,22] – thus it would be common to have ≥4 blood meals per lifetime [23]. Molecular analyses of bloodmeal DNA confirm multiple host feeds on capture [21,22,24,25]. In fact, re-feeding in the lab accelerates oocyst growth [18,19] and sporozoite development time, thereby shortening the incubation period and potentially increasing transmission risk [18].

With population-level drug deployment, infected mosquitoes would frequently be exposed to antimalarial-containing bloodmeals. Evidence from others supports that mosquitoes can absorb drugs during feeding. Duthaler, et al. demonstrated that the endectocide ivermectin can be absorbed in *Aedes* mosquitoes [26]. However, the question as to whether exposure to antimalarials within bloodmeals could significantly impact parasite development during sporogony has largely been unexplored. A seminal study on antimalarial activity across parasite life cycle stages supported potential anti-oocyst activity of several current antimalarials [27]. Although these data were indirectly inferred via eventual oocyst counts following drug exposure in gametocyte cultures, results support the theoretical potential for drugs to act during sporogonic stages. The authors reported that AQ, sulfamethoxazole, PYR, lumefantrine, mefloquine, pyronaridine, and an 8-aminoquinoline analog NPC1161B have potential activity against either gametocyte/exflagellation, ookinete, or oocyst stages [27]. Others have also reported incorporating drug exposure into infectious feeds can impact resultant mosquito oocysts and sporozoites [28–31], although much of this work utilized antibiotics or antimalarials not commonly used for mass prevention and treatment. Specific attention to antimalarials in widespread use and deployed with the goal of maximizing population coverage has not been examined.

To address this, we first investigated whether bloodmeal exposure to AQ, SP, and PPQ have direct impacts upon the *Anopheles* malaria vector at levels expected to be present in human blood. If so, such drug exposure would complicate further exploration of the effect of ingested antimalarials on parasite development within the mosquito, since the parasites require mosquitoes to survive for at least 9 days to complete sporogony and be transmitted onward. We focused on AQ/desethylamodiaquine (DEAQ), SP, and PPQ because of their relevance in prophylaxis and treatment, long half-lives, and literature supporting possible vector-stage activity [27]. We next examined whether these key chemoprophylactic antimalarials could be absorbed beyond the mosquito midgut, as such data would support the potential for them to act on parasites developing within the mosquito.

## Results

To investigate effects of mosquito ingestion of commonly used long-acting human antimalarials, we spiked drug into mosquito bloodmeals. We focused on AQ and DEAQ, the longer-acting metabolite of AQ that has potent anti-parasitic activity [32,33], as well as PPQ and SP. We used ivermectin (IVM) as a control due to its endectocidal properties [34] and related trials of mass distribution for malaria control [35–39]. The PK of these drugs are well characterized in humans by our group and others [33,40–46], informing relevant exposure concentrations in experiments performed here.

### Ingestion of drug solvents did not impact vector fitness of *Anopheles* mosquitoes

We first examined whether the solvents used to dissolve drug compounds had any impact on mosquito fitness at concentrations used with subsequent planned drug-spiked blood feeds. Solvents included acetonitrile (ACN):H_2_O 1:9 v/v with 0.5% formic acid (for AQ, DEAQ, PPQ), ACN:H_2_O 1:1 v/v with 0.1% hydrochloric acid (SDX), ACN:H2O 1:1 v/v/ with 0.5% formic acid (PYR), and DMSO (IVM). We calculated a maximum possible solvent exposure to involve a 1:833 plasma dilution, based on a modest drug dissolving target of 1 mg/ml, a standard example C_max_ of 300 ng/ml, plans to use drug concentrations up to twice the reported C_max_, and conservative estimates of pipetting variability. Feeding our lab colony *Anopheles gambiae* mosquitoes solvent diluted into plasma at 1:833 combined with packed RBCs to 40% hematocrit did not affect mosquito feeding behavior, survival, or fertility globally (S1 Fig and S1-S5 Tables). Survival curves were significantly different when comparing a few conditions (ACN 1:1 + 0.1% HCl lower survival than ACN 1:1 (p=0.01) and ACN 1:1 + FA 0.5% (p=0.0046); and HCl 0.1% alone lower survival than ACN 1:1 (p=0.0032), ACN 1:1 + FA 0.5% (p=0.0014), and FA 0.5% (p=0.0381)) but no solvent condition trialed was significantly different from the un-spiked blood only control (S4 Table). Subsequent drug stocks were made at 10 mg/ml for each compound, further reducing solvent concentration in drug-spiked bloodmeals. Given these results, DMSO (solvent for IVM) was not tested as a separate solvent control condition in subsequent experiments for lab mosquitoes.

### Ingestion of antimalarials AQ, DEAQ, PPQ, and SP has no significant impact on multiple measures of long-term lab-adapted *Anopheles gambiae* mosquito fitness

The malaria parasite’s ability to complete sporogony is significantly impacted by mosquito behavior and meal seeking, nutrient expenditure on ovulation, and overall vector lifespan [47]. To assess these effects, we next investigated the impact of drug ingestion on *Anopheles gambiae* mosquitoes from our laboratory colony. Drugs stocks of 10 mg/ml were diluted into plasma and combined with packed RBCs at 40% hematocrit for mosquito feeds. Target drug concentrations were selected based on literature reported C_max_ values following standard dosing (C_max_ approximated as 20 ng/ml for AQ [42], 500 ng/ml for DEAQ [42], 300 ng/ml for PPQ [41,43,44], 100 μg/ml for SDX [40,45], 400 ng/ml for PYR [40,45], 50 ng/ml for IVM [46]). Each drug was tested at concentrations ranging from one-half to twice C_max_. For SDX and PYR, given these components are dosed together, the SP combination was also tested. Negative controls included un-spiked blood and blood spiked with the relevant solvent at the equivalent volume and concentration. In addition, IVM served as a positive control given expected endectocidal activity upon ingestion.

Spiking drug into bloodmeals did not produce significant differences in mosquito feeding behavior when comparing control un-spiked blood feeds to each drug-spiked bloodmeal condition, as measured by percentage of female mosquitoes offered bloodmeal that fed (blood visible in abdomen) (Fig 1 A-C and S1 Table). Using an alternative chi-squared based analysis of fed versus unfed mosquitoes under each condition yielded some statistical differences. However, these differences were not consistent between drug concentrations nor even within the same control conditions (e.g. IVM across experimental groups) (S1 Supporting text and S2 Table).

**Fig 1.**
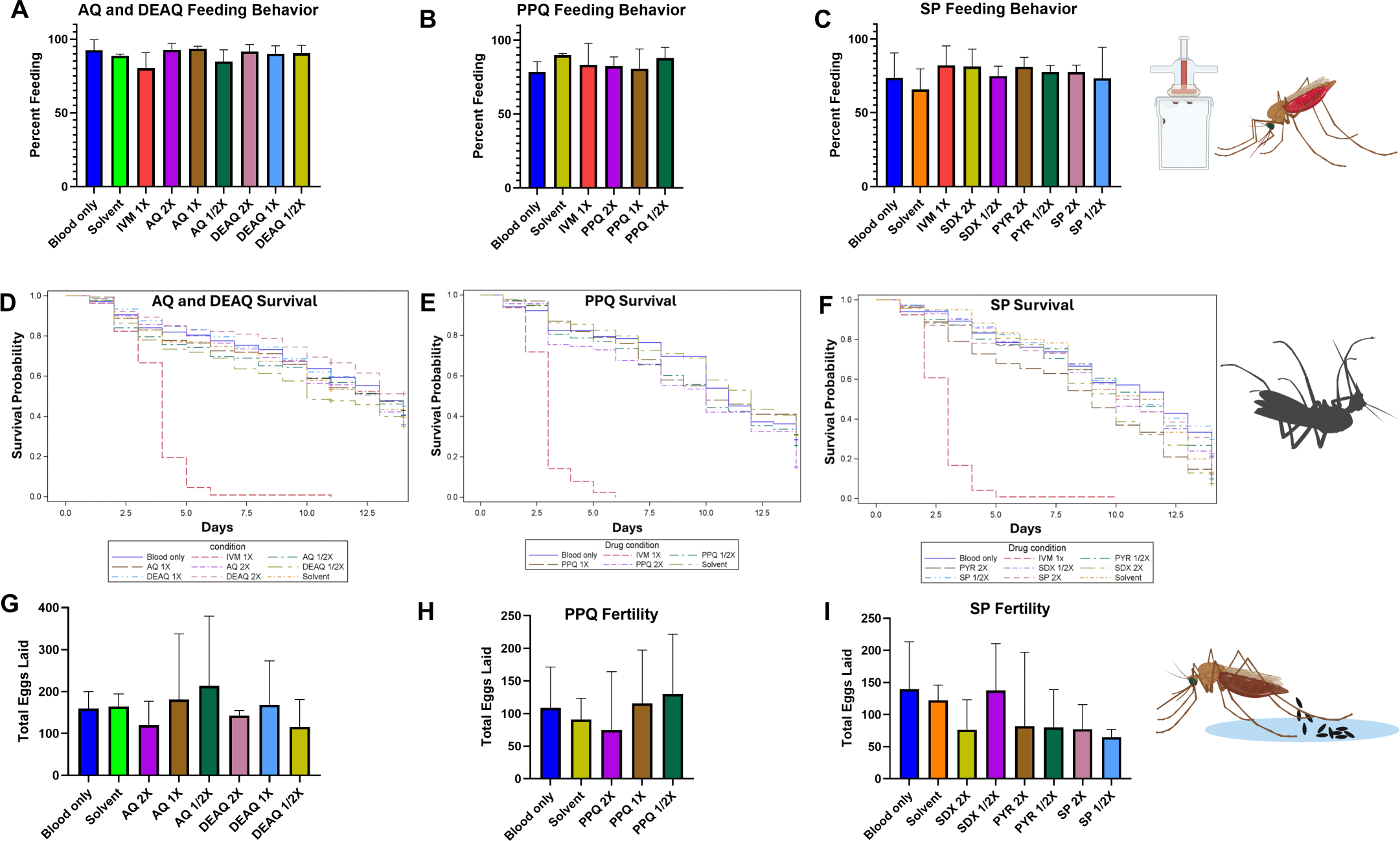
Tested antimalarials do not significantly impact lab-adapted *An. Gambiae* behavior, viability, or fertility. Human plasma was spiked with antimalarials AQ, DEAQ, PPQ, or SP at levels ranging from one half (1/2X) to twice (2X) the average reported C_max_ of standard human dosing, or with the endectocide IVM (at 1X C_max_), and combined with RBCs at 40% hematocrit for mosquito feeds. Negative controls included un-spiked blood (“Blood only”) or blood spiked with drug solvent alone (“Solvent”). Solvents were: ACN:H2O 1:9 + 0.5% FA for AQ, DEAQ, PPQ; ACN 1:1 H2O + 0.1% HCl for SDX; ACN 1:1 H2O + 0.5% FA for PYR (for SP experiments, “Solvent” represents the combination of both SDX and PYR solvents). Approximately 50 mosquitoes were fed per group. A-C) Mosquito feeding behavior plotted as percent taking a bloodmeal. D-F) Mosquito survival post drug ingestion plotted in a Kaplan-Meier survival curve (monitored for 14 days). G-I) Mosquito fertility assessed by total eggs laid per condition (n=5 mosquitoes monitored per experiment). Each graph represents combined data from 3 independent experiments (with 3 data points per condition/time, except for a few data points for survival monitoring resulting in fewer than triplicate observations for AQ and DEAQ Day 12-14, SP Day 9, 10, and PPQ D9 (2 data points each); and PPQ D 11, 13 (1 data point each)). In A-C and G-I, means are plotted with error bars representing SD.

Fed mosquitoes were monitored up to 14 days, towards the longer end of average expected duration of *P. falciparum* sporogony [17]. In activity monitoring experiments, consistent differences in behavior were not observed for any experimental condition other than following IVM feeds, which reliably led to loss of mobility notable within ∼24 hours (data not graphed). While survival rates naturally vary by individual experimental blood feed, particularly depending on age of mosquitoes at time of feed (ranging from 5-10 days), we also did not observe consistent negative impacts of drug ingestion on mosquito survival compared to control un-spiked or solvent-spiked bloodmeals other than expected effects from IVM (Fig 1 D-F, S1 Supporting text, and S3 Table). (Amongst multiple comparison analyses of all conditions, there were very limited conditions where survival curves were significantly different (S1 Supporting text, S4 Table).) Finally, there were also no significant differences in fertility measures following ingestion of any of the compounds compared to unaltered blood as indicated both by total egg counts per mosquitoes monitored (Fig 1 G-I; S5-S6 Tables) and egg counts per individual mosquitoes or percent of mosquitoes laying eggs (S2 Fig; S5-S6 Tables).

In contrast, IVM control feeds were consistently lethal, confirming successful target drug dilution precision and relevance of this experimental drug-spiked blood feeding system, Notably, while feeding behavior was unaltered in the presence of IVM, within 24 hours, the majority of mosquitoes lost mobility with significant numbers dying by 48 hours (see below and S3 Table). Because of this rapid alteration in behavior, IVM-fed mosquitoes were not subjected to individual fertility monitoring (otherwise drowning in forced egg laying conditions). When IVM-fed mosquitoes were allowed access to a communal egg laying water cup, they did not lay eggs (though dissections indicated eggs had formed, suggesting the reason for fertility absence was likely due to the neurologic and behavioral impacts of IVM). This is consistent with the known mechanism of action of IVM being a neurotoxin that binds to invertebrate-specific glutamate-gated chloride channels, causing hyperpolarization and paralysis [48].

### Ingestion of long-acting antimalarials did not significantly impact mosquito fitness of *Anopheles coluzzii* mosquitoes in Burkina Faso

Over many generations of lab adaptation, mosquito behavior and metabolism could be altered in ways that affect the impact of drug ingestion on fitness. Thus, we next investigated antimalarial feeding in field-derived mosquitoes from a routinely outbred *Anopheles coluzzii* colony in Burkina Faso. The same experimental design was employed, although the blood source differed, dissolving drug into commercial AB serum before mixing with donor packed RBCs from a community blood bank at 40% hematocrit. Field mosquito feeding rates using membrane feeders and stored blood components were significantly lower than for lab-adapted *Anopheles gambiae* tested at the Yale insectary (mean 51.6% of field mosquitoes feeding versus 84.1% of lab mosquitoes across all feeding experiments and conditions, p<0.0001), even with pre-feeding starvation for 24hrs. However, feeding rates did not differ by condition (percentage of female mosquitoes offered bloodmeal that fed, with blood visible in abdomen (Fig 2 A-C; S1 Table)). As above, using an alternative chi-squared based analysis of fed versus unfed mosquitoes under each condition yielded some statistical differences comparing mosquitoes fed spiked bloodmeals to those fed an unaltered bloodmeal or relevant solvent-spiked bloodmeal (S1 Supporting text, S2 Table). However, similar to the *An. gambiae* feeds, these differences were not consistent between drug concentrations nor even within the same control conditions across different feeding groups.

**Fig 2.**
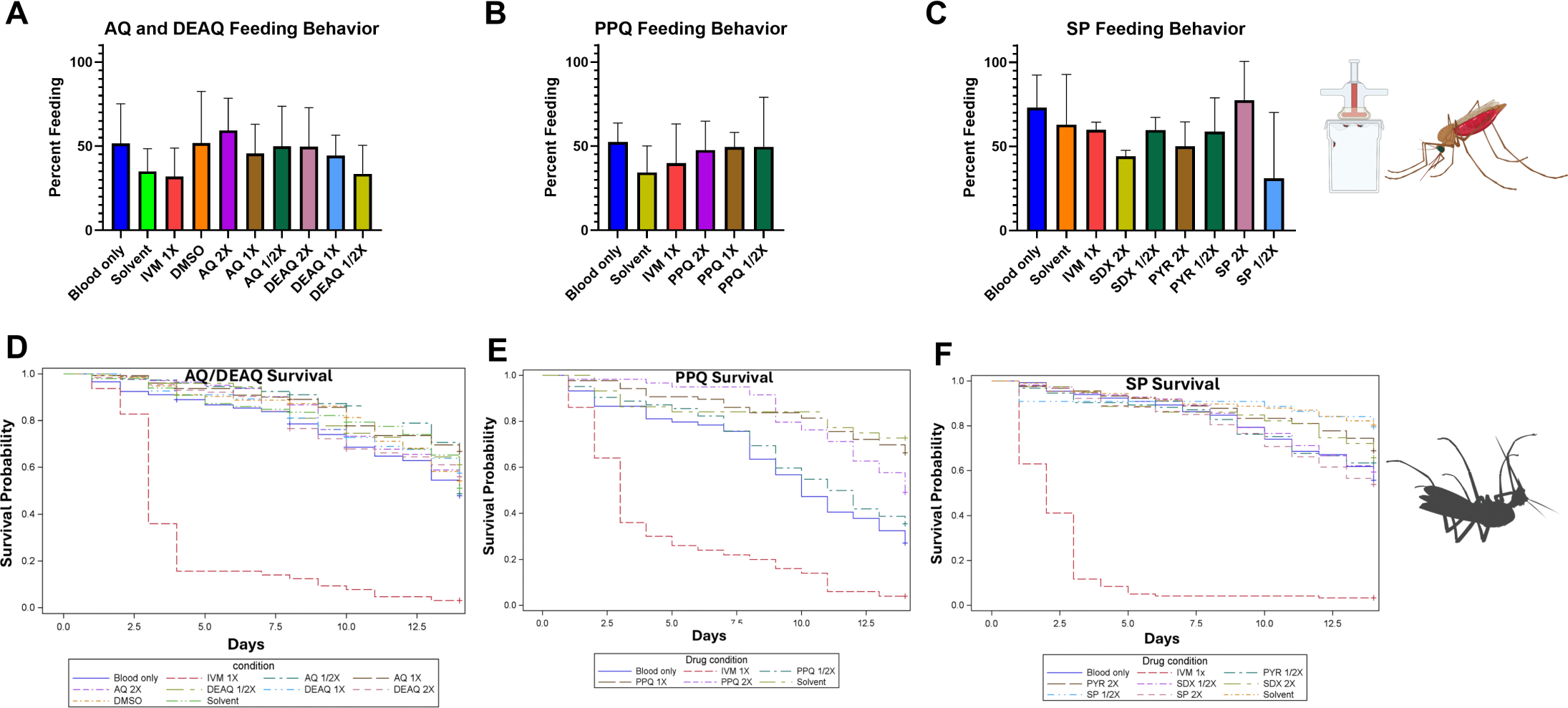
Tested antimalarials do not significantly impact field *An. coluzzii* behavior or viability. Human serum was spiked with antimalarials AQ, DEAQ, PPQ, or SP at levels ranging from one half (1/2X) to twice (2X) the average reported C_max_ of standard human dosing, or with the endectocide IVM (at 1X C_max_), and combined with RBCs at 40% hematocrit for mosquito feeds. Negative controls included un-spiked blood (“Blood only”) or that spiked with drug solvent alone (“Solvent”). Solvents were: ACN:H2O 1:9 + 0.5% FA for AQ, DEAQ, PPQ; ACN 1:1 H2O + 0.1% HCl for SDX; ACN 1:1 H2O + 0.5% FA for PYR (for SP experiments, “Solvent” represents the combination of both SDX and PYR solvents). DMSO (as solvent control for IVM drug) was also tested in AQ and DEAQ feeding experiments. Approximately 50 mosquitoes were fed per group. A-C) Mosquito feeding behavior plotted as percent taking a bloodmeal as percent taking a bloodmeal. Means are plotted with error bars representing SD. D-F) Mosquito survival post drug ingestion plotted in a Kaplan-Meier survival curve (monitored for 14 days). Each graph represents combined data from 3 independent experiments (with 3 data points per condition/time), unless otherwise specified: For AQ and DEAQ feeding and survival, experiments were repeated 4 times, but there were missing data points amounting to only triplicate survival observations for Day 5, 9, 11. For one SP experimental replicate, no mosquitoes fed for the SP1/2X condition (hypothesized due to physical feeder positioning), thus there were only duplicate observations for SP1/2X survival Day 1-14.

Nor were there consistent differences between control un-spiked versus drug-spiked blood feeds for any compound regarding routine daily behavior and survival, other than for IVM (Fig 2 D-F; S1 Supporting text, S3-S4 Tables). DMSO solvent was also confirmed to have no impact on field mosquito fitness measures when included as a separate control parameter (S3-S4 Tables). Between IVM feeds for all field mosquito experiments, mean 63.0% remained alive on D2 (SD 28.8), 32.1% on D3 (SD 27.7), and 15.8% on D4 (SD 23.4). These overall mortality rates were similar to lab-adapted *An. gambiae* IVM data (mean 70.8% alive on D2 (SD 29.7), 31.5% on D3 (SD 28.8), and 10.6% on D4 (SD 10.4)). However, for the *An. coluzzii* field mosquitoes, mean 11.5% remained alive by D7 (SD 21.9) and 3.1% by D14 (SD 4.9) after IVM feed, whereas for the *An. gambiae* lab mosquitoes, <1% remained alive by D6 after IVM feed (D6 mean 0.5% alive (SD 0.9)) (S3 Fig, S3 Table). Differences in overall survival curves comparing IVM-fed *An. gambiae* and *An. coluzzii* mosquitoes were also statistically significant (p<0.0001) (S3 Fig, S4 Table). For fertility assessments, field mosquitoes did not routinely lay eggs during each experimental replicate. However, these egg laying deficits also manifested following control un-spiked blood feeds. Of limited available data, no fertility differences were noted upon ingestion of any drug in terms of number of eggs laid per total mosquitoes, nor percent of mosquitoes laying eggs or number of eggs per laying mosquito (S4 Fig and S5-S6 Tables).

### DEAQ is absorbed after ingestion and distributed into *Anopheles gambiae* circulation and tissues

We hypothesized that ingested antimalarials could be absorbed from the midgut and circulate more widely, demonstrating the potential of drug exposure to later parasite stages in development. As a model system to assess this, we assayed the distribution of DEAQ in the mosquito using liquid chromatography with tandem mass spectrometry (LC-MS/MS) following drug-spiked blood feeds. Drug feeding conditions included DEAQ at one-half and twice C_max_ (of 500 ng/ml) as well as IVM at C_max_ (50 ng/ml), with mosquito tissue harvest occurring at 24 hrs and 120 hrs post feed (analyzing drug presence before and after completion of bloodmeal digestion at ∼72 hrs). We compared concentrations in 3 sample types: 1) whole mosquitoes, 2) midguts only, and 3) harvested hemolymph. We hypothesized midguts would contain the majority of drug following a bloodmeal, and that any drug presence in hemolymph would be indicative of drug absorption across the midgut into mosquito circulation. Whole mosquitoes or other target compartments were pooled (n=5) and collected in triplicate for each condition, with the overall experiment replicated twice.

Hemolymph collected via capillary perfusion of the mosquito thorax with PBS was assayed directly, whereas whole mosquitoes and midguts were harvested into an extraction solvent [26] and pulverized prior to mass spectrometry analysis. DEAQ had a limit of quantification of 0.5 ng/ml and was detected in a dose-dependent manner based on feeding at 2X or 1/2X C_max_ at 24 hrs in whole mosquitoes (mean 42.4 ng/ml or 6.7 ng/ml, respectively, p<0.0001), midguts (mean 33.2 ng/ml or 3.2 ng/ml, respectively, p=0.0082), and also hemolymph (mean 0.51 ng/ml for 2X C_max_; undetected at 1/2X C_max_ condition). DEAQ was also dose dependently detected based on feeding at either 2X or 1/2X C_max_ concentration at 120 hours in all compartments: whole mosquitoes (mean 33.1 ng/ml or 5.7 ng/ml, respectively, p=0.0001), midguts (mean 4.4 ng/ml or 0.6 ng/ml, respectively), and hemolymph (mean 0.9 ng/ml or 0.2 ng/ml, respectively). Between 24 to 120 hrs, DEAQ drug levels decreased in midguts but increased in hemolymph (trends present for both fed concentrations), providing evidence of absorption beyond midgut epithelium and prolonged circulation within hemolymph (Fig 3, Table 1, S7-S8 Tables). IVM had a limit of quantification of 1 ng/ml and was not detected other than in a few samples (detected in all three whole mosquito replicates in one experiment at average of 4.8 ng/ml). However, despite lacking extensive demonstration of biochemical drug absorption for IVM, mosquito behavior changed drastically upon IVM feed resulting in minimal movement and flaccid paralysis, highlighting the technical difficulties in capturing drug detection in individual mosquitoes even when it is clearly present. Harvest at 120 hr was not conducted for IVM because mosquitoes died before that time point was reached.

**Fig 3.**
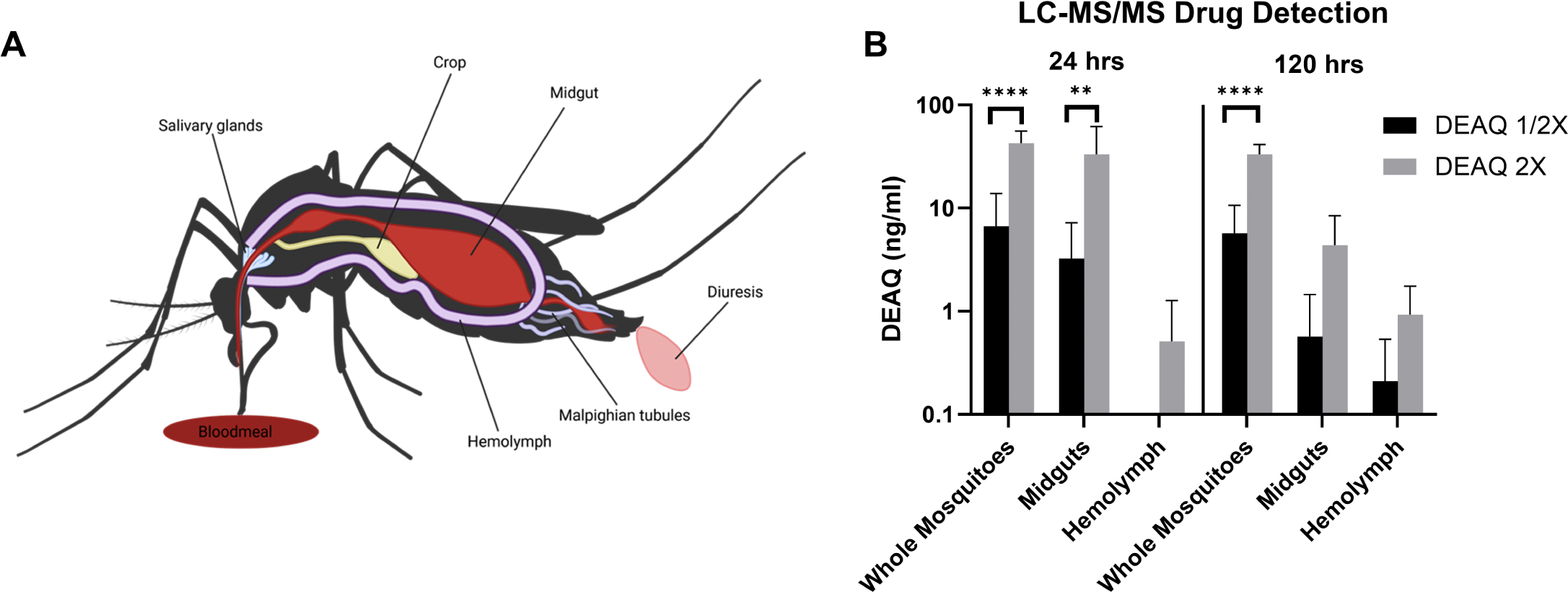
DEAQ is absorbed dose-dependently and detectable beyond midgut epithelium up to 5 days following bloodmeal ingestion. A) Cartoon depiction of mosquito anatomy upon bloodmeal ingestion. B) Mosquitoes were fed 1ɥg/ml or 250 ng/ml DEAQ to represent twice (2X) and half (1/2X) reported C_max_, respectively. Mosquitoes were harvested 24 hrs and 120 hrs post-feed, either collected whole or used for hemolymph collection and midgut dissection. For each condition, 5 of each mosquito sample type were pooled, with triplicates of pooled samples submitted to NorthEast BioLab (Hamden, CT) for LC-MS/MS analysis. Data represents average values from 2 separate experiments. Significant differences highlighted as **** = p<0.0001 and ** = p<0.01.

**Table 1.** DEAQ is absorbed and detectable in mosquito compartments up to 5 days beyond bloodmeal ingestion.

| **DEAQ detected in mosquitoes after spiked bloodmeal in ng/ml** |  |  |  |  |
| --- | --- | --- | --- | --- |
| **DEAQ fed concentration**<br>(relative to reported $C_{max}$) | **24h post feed** | | **120h post feed** | |
|  | **2X** | **1/2X** | **2X** | **1/2X** |
| **Whole mosquitoes** | 42.42<br>(13.48) | 6.67<br>(7.17) | 33.14<br>(8.30) | 5.71<br>(4.91) |
| **Midgut alone** | 33.22<br>(28.79) | 3.24<br>(3.96) | 4.37<br>(4.06) | 0.57<br>(0.88) |
| **Hemolymph alone** | 0.51<br>(0.76) | 0.00<br>(0.00) | 0.93<br>(0.83) | 0.21<sup>a</sup><br>(0.33) |

<sup>a</sup>The mean reported drug concentration at 120 hours for 1/2 $C_{max}$ condition was lower than the LOQ for hemolymph conditions due to some replicates with no detectable drug.

## Discussion

*Anopheles* mosquitoes, the vector for malaria transmission, are widely exposed to antimalarials by virtue of feeding on human hosts with long-acting drugs circulating in their blood. We fed *Plasmodium*-uninfected mosquitoes drug-spiked bloodmeals mimicking real-world drug concentrations of the antimalarials AQ, DEAQ, PPQ, and SP. We initially investigated uninfected mosquitoes to determine if drugs had any impact on mosquito viability which could in turn impact parasite transmission dynamics. We did not identify any consistent significant effects of these antimalarials on mosquito feeding behavior, activity levels, fertility, or survival up to 14 days following a drug-spiked blood meal (the time scale required to complete sporogony) in either a lab-adapted *Anopheles gambiae* mosquito strain or a field-maintained *Anopheles coluzzii* strain. Using DEAQ as a model due to its widespread use in SMC, we next assessed whether we could demonstrate absorption of long-acting antimalarials via blood-feeds using LC-MS/MS. We were able to detect drug present in whole mosquitoes, as well as within isolated midguts and hemolymph, 5 days post ingestion in a dose-dependent manner. The fact that drug levels decreased over time in midguts, while increasing in hemolymph, further supports that drugs present in mosquito bloodmeals can be absorbed beyond the midgut into the mosquito circulation and remain present in the circulation days after bloodmeals have been digested. To our knowledge, this is the first report of human antimalarial biochemical quantitative detection within *Anopheles* mosquito tissues following bloodmeal exposure.

With evidence that human antimalarials can be absorbed and persist in mosquito hemolymph, we hypothesize that drugs present in the bloodmeal have the potential to impact malaria parasite development and resistance selection during sporogony. Antimalarial resistance potentiation during mosquito stages has largely been overlooked compared to studies focused on the human blood stage. One area that has been examined is whether certain antibiotics can impact parasite development in the mosquito when ingested via blood meal [30,31]. The authors investigated penicillin-streptomycin, doxycycline, azithromycin, and trimethoprim-sulfamethoxazole. In some instances, antibiotics in feeds enhanced susceptibility of *An. gambiae* to *P. falciparum* infection (antibiotic exposure occurring at time of mosquito infection) and positively impacted vector lifespan and fecundity (postulated to be from mosquito gut microbiota disturbances, or possibly reduced levels of mosquito innate immune activation post blood feed) [30], though effects varied by antibiotic [31]. In particular, trimethoprim-sulfamethoxazole, an anti-folate similar to SP, reduced *P. falciparum* oocyst intensity. Recent work also has shown exposing mosquitoes to atovaquone starting at 48 hrs post-infection (via spiked sugar meal) significantly reduces oocyst size and sporozoite counts [29].

Antimalarial resistance develops primarily during symptomatic human blood-stage infection when replication rates and drug pressure are highest [11]. However, mosquito-level drug exposure could still impact already-developing parasites and contribute to strain selection at different points during sporogony. Studies investigating bottlenecks during sporogony emphasize the relevance of parasite oocyst and sporozoite numbers on overall transmission success, yet the specific determinants that impact parasite progression through vector-stage bottlenecks remain poorly understood [49,50]. Some research assessing infection of mosquitoes fed on gametocyte carriers with multiclonal infections has suggested both preferential transmission of minority clones to mosquitoes and propagation of minority clones through the mosquito, with significant implications for potential onward transmission of minority alleles including those that confer drug resistance [51,52].

While the DEAQ concentration we detected was low, our methods would underestimate drug levels given significantly diluted analyte and limited sampling times. The reported DEAQ IC_50_ of ∼60 nM [53,54] (∼24 ng/ml) is on par with levels we detected in whole mosquitoes and midguts at 24 hrs and whole mosquitoes at 120 hrs. Oocysts, developing between the mosquito midgut epithelium and basal lamina [55], could conceivably be exposed to ingested drug either directly via midgut bloodmeal contents (noting additional feeds stress and damage the midgut epithelial layer [21]) or via drug absorption into hemolymph (and subsequent tissue distribution). While oocysts are often portrayed as relatively “walled off” and thus potentially more protected from drug activity, atovaquone studies show that oocyst development can be impacted by either tarsal surface exposure (tissue absorption) or feeding [29]. Furthermore, drug present in midgut or circulation could impact pre-oocyst stages or sporozoites that travel via hemolymph to the salivary glands. We offer strong proof-of-concept data that support extensive drug distribution in mosquitoes for prolonged periods. Given 30-50% of some communities may be taking long-acting antimalarials, even rare selection events or low-level drug activity in mosquitoes could have large implications for malaria transmission dynamics in endemic areas.

There are several limitations to our present study. Although we only tested two *Plasmodium*-susceptible *Anopheles* species, we did compare lab and field mosquitoes, providing further evidence that the three antimalarials studied do not have an impact on mosquito fitness and viability. The IVM survival data highlights the relevance of testing field mosquitoes, given some were able to resist the endectocidal effect of IVM control feeds compared to lab mosquitoes (where <1% remained alive by D6). Future work would be bolstered by increasing the replicates/sample sizes from our field study, as we were not able to collect more fertility data given that mosquitoes did not lay eggs following feeds from some blood bank samples. We suspect this may be due to stored blood bank products suffering from breakdown of nutrients required for egg development. Despite this limitation, when egg laying did result from a feed, our data were consistent between control un-spiked vs. spiked drug feeds.

While some statistically significant differences in feeding behavior were present in alternative individual chi-squared analyses between individual spiked bloodmeal conditions in comparison to un-spiked blood or matching solvent-spiked blood, these differences were not consistent in direction between drug concentrations, mosquito species, nor between the same control conditions (e.g. IVM feeds) across different experimental groups. Thus, we believe they are more reflective of general variability between bloodmeal ingestion during any given experiment – particularly with the field mosquitoes that are less adapted to artificial membrane feeding. Similarly, isolated survival curves with significant differences are likely more reflective of one-off technically significant results that arise in the myriad multiple comparisons done within each drug or solvent feeding group and mosquito species, as there was no consistency in significance across drug concentration and mosquito species. Minimally, our results show that drug presence did not negatively impact survival, with most of the significant curve differences showing the lower concentration of drug or drug-absent blood only condition had reduced survival.

We focused our initial drug detection analyses on DEAQ given its widespread use in both treatment and prevention, with the primary goals of determining whether we could 1) detect drug in various mosquito compartments, and 2) observe any concentration changes over time. Our time- and labor-intensive design was successful in achieving these objectives, and we are not surprised that there was variability in terms of the number of samples with detectable drug, particularly since concentrations were often near the limit of LC-MS/MS quantification. This is true for IVM, which was undetectable in our study with a LOQ of 1 ng/ml. However, IVM was clearly absorbed, as evidenced by mosquito death following ingestion. Given these experimental limitations, the fact that we could successfully detect DEAQ in individual mosquitoes days after ingestion, even in isolated hemolymph, supports that antimalarial absorption occurs, and can potentially interface with different stages of sporogonic development. Furthermore, our DEAQ detection results were highly specific (with no drug detection in any negative controls fed solvent-spiked blood or unaltered blood) and concentration dependent (in terms of initial fed drug amount).

Having demonstrated mosquito drug exposure occurs but does not itself negatively impact the vector, the next question will be investigating the impact of antimalarial exposure on infected mosquito oocyst and sporozoite output, as well as assessing whether drugs could impact the success of strains developing in the mosquito, particularly in mixed infections where less resistant strains may have a developmental advantage. Given rapidly expanding drug resistance to artemisinin based antimalarials now in Africa [4], concern about partner drug resistance that could contribute to failure of first-line treatment, and stagnating progress on reducing the global malaria burden in recent years, the malaria community needs to consider all avenues of possible contribution to drug-resistance spread. This includes the possibility that human antimalarials could impact vector stage transmission dynamics. Better understanding of mosquito drug exposure and its impact could also help in developing strategies to target the parasite within the mosquito.

## Materials and methods

### Mosquito colonies

Two species of mosquitoes were used in this study: 1) *An. gambiae* “G3” strain, a highly lab-adapted, insecticide-susceptible strain of African origin (obtained from the Catteruccia lab, Harvard University) reared at Yale lab facilities in New Haven, CT under standard conditions (27 ± 2°C, 70% humidity, 12 hour light cycle with access to 10% glucose solution); referred to as “lab mosquitoes;” and 2) *Anopheles coluzzii* outbred mosquitoes (established in 2023 and repeatedly replenished with F1 from wild-caught female mosquitoes collected in Bama, Burkina Faso (coordinates 12.0395° N, 4.4010° W; ∼60km north of Bobo-Dioulasso); field-collected female mosquitoes morphologically identified as belonging to the *An. gambiae* complex further identified by PCR before pooling) and maintained either in the Institut de Recherche en Sciences de la Santé (IRSS) laboratory in Bobo Dioulasso under standard conditions (27 ± 2°C, 70% humidity, 12 hour light cycle with access to 10% glucose solution) or in the Bama malaria sphere under semi-natural conditions (exposed to all environmental factors including temperature and relative humidity). These mosquitoes (maintained in either Bama malaria sphere or Bobo-Dioulasso lab conditions) are collectively referred to as “field mosquitoes.”

### Drug-spiked blood feeds

Fresh human blood was routinely obtained from healthy O+ donors in New Haven, CT or via hospital blood bank in Bobo-Dioulasso, Burkina Faso. For human donors at Yale, plasma was removed and packed RBCs were separated and washed with un-supplemented RPMI media. Blood products in Burkina Faso were obtained as packed RBCs and used as is. Drugs of interest were dissolved into relevant solvent at 10mg/ml and stored at -20°C. Briefly, amodiaquine (AQ; Sigma), desethylamodiaquine (DEAQ; Cayman), and piperaquine (PPQ; Sigma) were dissolved into 1:9 v/v acetonitrile:H2O with 0.5% formic acid; sulfadoxine (SDX; Cayman) was dissolved into 1:1 v/v acetonitrile:H2O with 0.1% hydrochloric acid; pyrimethamine (PYR: Cayman) was dissolved into 1:1 v/v acetonitrile:H2O with 0.5% formic acid; and ivermectin (IVM; Sigma) was dissolved into DMSO. Drug stocks were diluted into either plasma (Yale donors from above) or commercial non-immune AB serum (Sigma; used for Burkinabé experiments) to achieve physiologic concentrations in the range of one-half to two times the average reported C_max_ after a standard human oral dose as per available literature (C_max_ approximated as 20 ng/ml for AQ [42], 500 ng/ml for DEAQ [42], 300 ng/ml for PPQ [41,43,44], 100 ɥg/ml for SDX [40,45], 400 ng/ml for PYR [40,45], 50 ng/ml for IVM [46]). Drug-spiked plasma/serum was then combined with packed RBCs at 40% hematocrit and fed for up to 60 minutes to 5-10 day old *Anopheles* mosquitoes via standard membrane feeding practice with pre-warmed glass feeders and parafilm membrane.

### Mosquito viability assessments

50 female mosquitoes per condition were placed in individual containers for drug-spiked blood feeds, after which fed mosquitoes were manually separated and maintained at standard conditions. For all experiments, mosquitoes were only offered one singular feeding opportunity and never refed. Feeding behavior was determined by the percentage of female mosquitoes present that imbibed a bloodmeal. Post feeding, viability was monitored for each condition for up to 14 days, to span the typical amount of time sporozoites develop in a mosquito. Daily mosquito behavior was also monitored by gently blowing on the cartons of mosquitoes and assessing flight activity. Finally, 2 days after feeding, 5 randomly selected individual mosquitoes per condition were placed in individual containers for forced egg laying, with the number of eggs per mosquito recorded, as well as the number of laying mosquitoes. (Fertility assessments did not take place for the ivermectin condition because by this time post feed, these mosquitoes were notably unhealthy and not able to withstand forced egg laying conditions.) Experiments were repeated in triplicate. Statistical comparisons between mosquito fertility were assessed via ordinary one-way ANOVA with Dunnett’s multiple comparisons test to compare each condition to “blood only” control (performed in GraphPad Prism version 10.5.0). Statistical comparisons between mosquito feeding behavior were assessed both via ordinary one-way ANOVA of percent of mosquitoes feeding with Dunnett’s multiple comparisons test to compare each condition to “blood only” or control, as well as using contingency table analyses of total fed and unfed mosquitoes combining the data from multiple experiments for each drug feeding group and comparing a given experimental condition to the control “blood only” or “solvent” condition (both performed in GraphPad Prism version 10.5.0). Survival was assessed via stratified log-rank test of Kaplan Meier analysis combining the data from multiple experiments for each drug feeding group and using the Tukey-Kramer adjustment for multiple comparisons (performed in SAS 9.4, Cary, NC).

### Biochemical drug detection in the mosquito

We analyzed drug absorption within individual mosquitoes using IVM (at 24 hours post feed) and DEAQ (at 24 hours and 120 hours post feed). 120 mosquitoes were fed per condition (control blood only, IVM 1X, ACN:H_2_O 1:9 + 0.5% FA solvent, DMSO solvent, DEAQ 2X, DEAQ 1/2X). Fed mosquitoes were manually separated and maintained at standard conditions. Per condition, 5 mosquitoes were pooled into 100 µl “extraction solvent” (70% MeOH + 0.1% formic acid, per others’ reported methods [26]), 5 mosquito midguts were harvested into 50 µl of extraction solvent, and hemolymph was pooled from 5 mosquitoes; these were all collected in triplicate. To collect hemolymph, mosquitoes were immobilized on ice under a dissecting microscope. The last segment of the abdomen was removed. A glass capillary tube made to fine needle point attached to 1 mm diameter silicone tubing and a 100 µl Hamilton syringe mounted on an automated microdispenser (PB600 Repeating Syringe Dispenser, Hamilton) filled with PBS was used to puncture the thorax underneath the wings, with 2 µl of PBS injected in increments. Resultant hemolymph expelled out of the dissected abdomen was collected via p10 pipette (approximately 2-5 µl of hemolymph diluted into PBS collected per mosquito). Hemolymph-PBS solutions were not further processed other than high speed centrifugation at 16.1g for 10 minutes with supernatant submitted for analysis. Whole mosquitoes and midguts were pulverized using the pellet pestle grinder (DWK Life Sciences), vortexed for 30 sec, sonicated for 10 min, and centrifuged at 16.1g for 10 min with supernatants submitted for LC-MS/MS drug detection (similar to described [26]). Drug detection was performed by NorthEast BioLabs (Hamden, CT) with appropriate standard calibration using control matrix consisting of sugar-fed whole mosquitoes in extraction solvent. Reported limits of quantification were 1 ng/ml for ivermectin and 0.5 ng/ml for desethylamodiaquine. Data represents results from triplicate pooled samples for each harvest parameter, averaged between two identical repeated experiments. Statistical comparisons between drug levels detected in different mosquito compartments were assessed via one-way ANOVA followed by pairwise comparisons of interest using Holm-Šídák’s multiple comparisons test (calculated using GraphPad Prism version 10.5.0).

### Ethics statement

Human subjects work involving use of blood from U.S. donors was approved by the Yale University Institutional Review Board (IRB 2000034487). Formal verbal consent was obtained for all subjects involved. Blood from Burkina Faso donors was obtained via local hospital blood bank in Bobo-Dioulasso, Burkina Faso (IRB exempt de-identified blood products).

## Acknowledgements

We would like to acknowledge the larger insectary team at IRSS in Burkina Faso (Hien Dombagniro Raymond, Somda Mouonyr Stéphane, Helène Somé, and Somda Gninéabéka Béranger Eustache) for helping rear mosquitoes and assist with experimental feeds. We would also like to acknowledge the team at NorthEast BioAnalytical Laboratories for their willingness to work with us to pursue drug detection analyses in a novel mosquito matrix. Finally, we would also like to thank all of the blood donors both at Yale and in Burkina Faso. Cartoon representations in all figures were created in BioRender (https://BioRender.com).

## Supporting information

### S1 Supporting text

**S1 Fig. Solvents do not significantly impact lab-adapted *An. gambiae* behavior, viability, or fertility.** Solvents used to dissolve drugs AQ, DEAQ, PPQ, SDX, PYR, and IVM were diluted into plasma at 1:833 dilution and combined with RBCs at 40% hematocrit for mosquito feeds. Un-spiked blood (“Blood only”) served as control. Abbreviations include: ACN – acetonitrile, used at either 1:1 or 1:9 v/v with H_2_O; FA – formic acid; HCl – hydrochloric acid. Approximately 50 mosquitoes were fed per group. A) Mosquito feeding behavior plotted as percent taking a bloodmeal. B) Mosquito survival post solvent ingestion plotted in a Kaplan-Meier survival curve (monitored for 14 days). C) Mosquito fertility assessed by total eggs laid per condition (all mosquitoes monitored communally per experiment). Data are from experiments performed in duplicate (except for only single survival data points available for D9 and D11 ACN:H_2_O 1:1 + HCl 0.1% and HCl 0.1% conditions). In A and C, means are plotted with error bars representing SD.

**S2 Fig. Tested antimalarials do not significantly impact percent of lab-adapted *An. gambiae* mosquitoes laying eggs.** Human plasma was spiked with antimalarials AQ, DEAQ, PPQ, or SP at levels ranging from one half (1/2X) to twice (2X) the average reported C_max_ of standard human dosing and combined with RBCs at 40% hematocrit for mosquito feeds. Negative controls included un-spiked blood (“Blood only”) or blood spiked with drug solvent alone (“Solvent”). Solvents were: ACN:H_2_O 1:9 + 0.5% FA for AQ, DEAQ, PPQ; ACN 1:1 H_2_O + 0.1% HCl for SDX; ACN 1:1 H_2_O + 0.5% FA for PYR (for SP experiments, “Solvent” represents the combination of both SDX and PYR solvents). Approximately 50 mosquitoes were fed per group, measuring effect of A) AQ and DEAQ, B) PPQ, and C) SP on mosquito fertility as measured by percent mosquitoes laying eggs (of n=5 mosquitoes monitored per experiment). Each graph represents combined data from 3 independent experiments. Means are plotted with error bars representing SD. There were no significant differences in percent of egg laying mosquitoes between groups.

**S3 Fig. Reduced IVM killing effects in field-derived *An. coluzzii* versus lab-derived *An. gambiae* mosquitoes.** Human plasma was spiked with the endectocide IVM (at 1X C_max_ of standard human dosing) and combined with RBCs at 40% hematocrit for mosquito feeds. Un-spiked blood (“Blood only”) served as control. Approximately 50 mosquitoes were fed per group. Mosquito survival post drug ingestion plotted in a Kaplan-Meier survival curve (monitored for 14 days). Data here is combined from multiple available independent feeding experiments (9 for *An. gambiae* and 10 for *An. coluzzii*, except the “Blood only” condition for *An. gambiae* had 8 data points for D 10, 11, 12, 14, and 6 data points for D9, 13; and both conditions for *An. coluzzii* had 9 data points for D 5, 9, 11). There were no significant differences in “Blood only” survival between lab and field mosquitoes, however for “IVM 1X” the *An. coluzzii* mosquitoes demonstrated significantly improved survival compared to *An. gambiae* mosquitoes (p<0.0001).

**S4 Fig. Tested antimalarials do not significantly impact field-adapted *An. coluzzii* fertility.** Human serum was spiked with antimalarials AQ, DEAQ, PPQ, or SP at levels ranging from one half (1/2X) to twice (2X) the average reported C_max_ of standard human dosing and combined with RBCs at 40% hematocrit for mosquito feeds. Negative controls included un-spiked blood (“Blood only”) or blood spiked with drug solvent alone (“Solvent”). Solvents were: ACN:H_2_O 1:9 + 0.5% FA for AQ, DEAQ, PPQ; ACN 1:1 H_2_O + 0.1% HCl for SDX; ACN 1:1 H_2_O + 0.5% FA for PYR (for SP experiments, “Solvent” represents the combination of both SDX and PYR solvents). Approximately 50 mosquitoes were fed per group. Mosquito fertility (of n=5 mosquitoes monitored per experiment) is depicted both by total eggs laid per condition (A-C) and percent of mosquitoes laying eggs (D-F). Each graph represents combined data from 3 independent experiments, however for many experimental replicates, mosquitoes did not routinely lay eggs – for AQ and DEAQ feeds, there was only egg laying data from 2 replicates (and the DEAQ 2X-fed group from one additional replicate); for PPQ feeds, there was only egg laying data from 1 replicate (and the PPQ 2X-fed group from one additional replicate). Means are plotted with error bars representing SD.

**S1 Table. Feeding rates for lab-derived *An. gambiae* and field-derived *An. coluzzii* offered bloodmeals spiked with solvents and tested antimalarials.** Data represents the percent of female mosquitoes that fed (abdomen with visible bloodmeal) under each condition, reported as the mean between experimental replicates (and Standard Deviation) (two replicates for initial solvent control feeding; three replicates for all drug feeding experiments). Plasma was spiked with solvents DMSO, ACN:H_2_O 1:1 v/v + 0.5% FA, ACN:H_2_O 1:9 v/v + 0.5% FA, ACN:H_2_O 1:1, FA 0.5%, ACN:H_2_O 1:1 + 0.1% HCl, and 0.1% HCl all used at 1:833 dilution in plasma and mixed with RBCs to achieve 40% hematocrit in initial control experiments using lab-derived *An. gambiae*. In further experiments with both lab *An. gambiae* and field *An. coluzzii*, plasma or serum was spiked with the antimalarials AQ, DEAQ, PPQ, or SP at levels ranging from one half (1/2X) to twice (2X) the average reported C_max_ of standard human dosing, or with the endectocide IVM (at 1X C_max_), and combined with RBCs at 40% hematocrit for mosquito feeds. Negative controls included un-spiked blood (“Blood only”) or for the drug feeding experiments, blood spiked with the relevant drug solvent alone (“Solvent”). Solvents used in the drug feeding experiments were: ACN:H_2_O 1:9 + 0.5% FA for AQ, DEAQ, PPQ; ACN:H_2_O 1:1 + 0.1% HCl for SDX; ACN:H_2_O 1:1 + 0.5% FA for PYR (for SP experiments, “Solvent” represents the combination of both SDX and PYR solvents). Approximately 50 mosquitoes were fed per group.

**S2 Table. Raw feeding data and contingency analyses for lab-derived *An. Gambiae* and field-derived *An. coluzzii* offered bloodmeals spiked with solvents and tested antimalarials.** Data represents the number of female mosquitoes that fed (abdomen with visible bloodmeal) or did not feed under each condition, reported as the total between experimental replicates (two replicates for initial solvent control feeding; three replicates for all drug feeding experiments). Contingency analyses comparing the rates of fed to unfed mosquitoes for each experimental condition are included against the blood only or relevant solvent controls, as well as for total lab An. gambiae and field An. coluzzii feeding rates under all conditions (reported as p values, italicized if <0.05 with direction of significance stated). Plasma was spiked with solvents DMSO, ACN:H_2_O 1:1 v/v + 0.5% FA, ACN:H_2_O 1:9 v/v + 0.5% FA, ACN:H_2_O 1:1, FA 0.5%, ACN:H_2_O 1:1 + 0.1% HCl, and 0.1% HCl all used at 1:833 dilution in plasma and mixed with RBCs to achieve 40% hematocrit in initial control experiments using lab-derived *An. gambiae*. In further experiments with both lab *An. gambiae* and field *An. coluzzii*, plasma was spiked with the antimalarials AQ, DEAQ, PPQ, or SP at levels ranging from one half (1/2X) to twice (2X) the average reported C_max_ of standard human dosing, or with the endectocide IVM (at 1X C_max_), and combined with RBCs at 40% hematocrit for mosquito feeds. Negative controls included un-spiked blood (“Blood only”) or for the drug feeding experiments, blood spiked with the relevant drug solvent alone (“Solvent”). Solvents used in the drug feeding experiments were: ACN:H_2_O 1:9 + 0.5% FA for AQ, DEAQ, PPQ; ACN:H_2_O 1:1 + 0.1% HCl for SDX; ACN:H_2_O 1:1 + 0.5% FA for PYR (for SP experiments, “Solvent” represents the combination of both SDX and PYR solvents). Approximately 50 mosquitoes were fed per group.

**S3 Table. Percent survival of lab-derived *An. gambiae* and field-derived *An. coluzzii* following ingestion of bloodmeal spiked with solvents and tested antimalarials.** Data represents percent survival of mosquitoes (approximately n=50 fed; unfed removed) over time after feeding on bloodmeals spiked with solvents and drugs of interest. Data are from 3 experimental replicates, with individual data for each time/condition shown, unless otherwise indicated (blank for few times/conditions when data was not captured; only 2 replicates for initial solvent control experiments; 4 replicates for *An. coluzzii* AQ/DEAQ feeds; 9 replicates combining all *An. gambiae* IVM feeding data and 10 replicates combining all *An. coluzzii* IVM feeding data). Means (and Standard Deviation) are also shown for each time/condition. Plasma was spiked with solvents DMSO, ACN:H_2_O 1:1 v/v + 0.5% FA, ACN:H_2_O 1:9 v/v + 0.5% FA, ACN:H_2_O 1:1, FA 0.5%, ACN:H_2_O 1:1 + 0.1% HCl, and 0.1% HCl all used at 1:833 dilution in plasma and mixed with RBCs to achieve 40% hematocrit in initial control experiments using lab-derived *An. gambiae*. In further experiments with both lab *An. gambiae* and field *An. coluzzii*, plasma was spiked with the antimalarials AQ, DEAQ, PPQ, or SP at levels ranging from one half (1/2X) to twice (2X) the average reported C_max_ of standard human dosing, or with the endectocide IVM (at 1X C_max_), and combined with RBCs at 40% hematocrit for mosquito feeds. Negative controls included un-spiked blood (“Blood only”) or for the drug feeding experiments, blood spiked with the relevant drug solvent alone (“Solvent”). Solvents used in the drug feeding experiments were: ACN:H_2_O 1:9 + 0.5% FA for AQ, DEAQ, PPQ; ACN:H_2_O 1:1 + 0.1% HCl for SDX; ACN:H_2_O 1:1 + 0.5% FA for PYR (for SP experiments, “Solvent” represents the combination of both SDX and PYR solvents).

**S4 Table. Survival curve comparisons for lab-derived *An. gambiae* and field-derived *An. coluzzii* following ingestion of bloodmeal spiked with solvents and tested antimalarials.** Data represents Kaplan-Meier survival curve comparisons between specified groups, with Chi square results and p values (unadjusted and adjusted for multiple comparisons using Tukey-Kramer post-hoc analyses). Significant adjusted p values <0.05 are italicized for comparisons not involving IVM, with the direction of significance noted. Survival curves for each group are from 3 experimental replicates (other than only 2 replicates for initial solvent control experiments; 4 replicates for An. coluzzii AQ/DEAQ feeds). Data comparing survival curves for all blood only controls and IVM 1X feeds in *An. gambiae* versus *An. coluzzii* mosquitoes is also included. Plasma was spiked with solvents DMSO, ACN:H_2_O 1:1 v/v + 0.5% FA, ACN:H_2_O 1:9 v/v + 0.5% FA, ACN:H_2_O 1:1, FA 0.5%, ACN:H_2_O 1:1 + 0.1% HCl, and 0.1% HCl all used at 1:833 dilution in plasma and mixed with RBCs to achieve 40% hematocrit in initial control experiments using lab-derived *An. gambiae*. In further experiments with both lab *An. gambiae* and field *An. coluzzii*, plasma was spiked with the antimalarials AQ, DEAQ, PPQ, or SP at levels ranging from one half (1/2X) to twice (2X) the average reported C_max_ of standard human dosing, or with the endectocide IVM (at 1X C_max_), and combined with RBCs at 40% hematocrit for mosquito feeds. Negative controls included un-spiked blood (“Blood only”) or for the drug feeding experiments, blood spiked with the relevant drug solvent alone (“Solvent”). Solvents used in the drug feeding experiments were: ACN:H_2_O 1:9 + 0.5% FA for AQ, DEAQ, PPQ; ACN:H_2_O 1:1 + 0.1% HCl for SDX; ACN:H_2_O 1:1 + 0.5% FA for PYR (for SP experiments, “Solvent” represents the combination of both SDX and PYR solvents).

**S5 Table. *An. gambiae* and *An. coluzzii* mosquito fertility following ingestion of bloodmeal spiked with solvents and tested antimalarials.** For initial solvent control feeds in *An. gambiae*, data includes the total number of eggs laid and eggs laid per mosquito (all fed mosquitoes monitored communally for egg laying; percent of mosquitoes laying eggs unknown) from two experimental replicates. For all further drug feeds, data includes the total eggs laid, number of eggs laid per mosquito, and percent of mosquitoes laying eggs (n=5 mosquitoes monitored in egg laying conditions) following bloodmeal ingestion under each condition between experimental replicates (three replicates for *An. gambiae* and *An. coluzzii* DEAQ and SP feeds; 2 replicates for *An. coluzzii* PPQ feeds excluding further replicates from analyses where no mosquitoes laid eggs). Data is reported as the mean (with Standard Deviation). Plasma was spiked with solvents DMSO, ACN:H_2_O 1:1 v/v + 0.5% FA, ACN:H_2_O 1:9 v/v + 0.5% FA, ACN:H_2_O 1:1, FA 0.5%, ACN:H_2_O 1:1 + 0.1% HCl, and 0.1% HCl all used at 1:833 dilution in plasma and mixed with RBCs to achieve 40% hematocrit in initial control experiments using lab-derived *An. gambiae*. In further experiments with both lab *An. gambiae* and field *An. coluzzii*, plasma was spiked with the antimalarials AQ, DEAQ, PPQ, or SP at levels ranging from one half (1/2X) to twice (2X) the average reported C_max_ of standard human dosing, and combined with RBCs at 40% hematocrit for mosquito feeds. Negative controls included un-spiked blood (“Blood only”) or for the drug feeding experiments, blood spiked with the relevant drug solvent alone (“Solvent”). Solvents used in the drug feeding experiments were: ACN:H_2_O 1:9 + 0.5% FA for AQ, DEAQ, PPQ; ACN:H_2_O 1:1 + 0.1% HCl for SDX; ACN:H_2_O 1:1 + 0.5% FA for PYR (for SP experiments, “Solvent” represents the combination of both SDX and PYR solvents).

**S6 Table. Lab-derived *An. gambiae* and field-derived *An. coluzzii* mosquito egg laying data following ingestion of bloodmeal spiked with tested antimalarials.** Data shown as the number of eggs laid per mosquito (n=5 mosquitoes monitored in individual egg laying conditions) following bloodmeal ingestion under each condition in three experimental replicates per group. Plasma was spiked with the antimalarials AQ, DEAQ, PPQ, or SP at levels ranging from one half (1/2X) to twice (2X) the average reported C_max_ of standard human dosing and combined with RBCs at 40% hematocrit for mosquito feeds. Negative controls included un-spiked blood (“Blood only”), or for the drug feeding experiments, blood spiked with the relevant drug solvent alone (“Solvent”) (DMSO also included as a separate solvent in *An. coluzzii* AQ/DEAQ feeds). Solvents used in the drug feeding experiments were: ACN:H_2_O 1:9 + 0.5% FA for AQ, DEAQ, PPQ; ACN:H_2_O 1:1 + 0.1% HCl for SDX; ACN:H_2_O 1:1 + 0.5% FA for PYR (for SP experiments, “Solvent” represents the combination of both SDX and PYR solvents).

**S7 Table. Drug detection in isolated *An. gambiae* mosquito compartments.** Mosquitoes were fed 1ɥg/ml or 250 ng/ml DEAQ to represent twice (2X) and half (1/2X) reported C_max_, respectively, or 50 ng/ml IVM to represent 1X reported Cmax. Controls included un-spiked blood (Blood only), and solvent-spiked blood (ACN:H_2_O 1:9 + 0.5% FA for DEAQ; DMSO for IVM). Mosquitoes were harvested 24 hrs (all conditions) and 120 hrs (Blood only, ACN+FA, DEAQ 2X, DEAQ 1/2X) post-feed, either collected whole or used for hemolymph collection and midgut dissection. For each condition, 5 of each mosquito sample type were pooled, with triplicates of pooled samples submitted to NorthEast BioLab (Hamden, CT) for LC-MS/MS analysis. All data from 2 separate experiments is shown. B=blank (no drug detected).

**S8 Table. Drug detection comparisons in isolated *An. gambiae* mosquito compartments over time.** Mosquitoes were fed 1 ɥg/ml or 250 ng/ml DEAQ to represent twice (2X) and half (1/2X) reported C_max_, respectively, or 50 ng/ml IVM to represent 1X reported C_max_. Controls included un-spiked blood (Blood only), and solvent-spiked blood (ACN:H_2_O 1:9 + 0.5% FA for DEAQ; DMSO for IVM). Mosquitoes were harvested 24 hrs (all conditions) and 120 hrs (Blood only, ACN:H_2_O 1:9 + 0.5% FA, DEAQ 2X, DEAQ 1/2X) post-feed, either collected whole or used for hemolymph collection and midgut dissection. For each condition, 5 of each mosquito sample type were pooled, with triplicates of pooled samples submitted to NorthEast BioLab (Hamden, CT) for LC-MS/MS analysis. Data is from 2 separate experiments. Results are listed for one-way ANOVA analyzing drug levels detected in whole mosquitoes, midguts, and hemolymph with pairwise comparisons of conditions of interest over time. Significant p values (p<0.05) are italicized.

