## Supporting text for "*Anopheles* mosquitoes exposed to long-acting antimalarials via drug-spiked bloodmeal absorb drug but do not suffer fitness costs"

### S1 Supplemental text

#### Additional statistical results descriptions for *An. gambiae* mosquito fitness tests

##### Feeding behavior

For mosquito feeding behavior experiments assessing the number of mosquitoes ingesting bloodmeals, beyond analyzing the percent of mosquitoes feeding under any condition, we additionally used chi-squared analysis to compare fed versus unfed mosquitoes for each condition. The chi-squared analyses identified some statistically significant differences comparing mosquitoes fed spiked bloodmeals to those fed an unaltered bloodmeal: mosquitoes within the AQ/DEAQ feeding group fed less with IVM 1X (p=0.0004) and AQ 1/2X (p=0.0104) spiked bloodmeals; within the PPQ feeding group fed more with ACN:$H_2O$ 1:9 + FA 0.5% solvent (p=0.006) and PPQ 1/2X (p=0.036); and within the SP feeding group fed more with IVM 1X (p=0.047). This chi-squared analyses also identified some statistically significant differences comparing mosquitoes fed spiked bloodmeals to those fed relevant solvent-spiked bloodmeals: mosquitoes within the AQ/DEAQ feeding group fed less with IVM 1X spiked bloodmeal (p=0.029); mosquitoes within the PPQ feeding group fed less for PPQ 1X (p=0.0203); mosquitoes within the SP feeding group fed more with IVM 1X (p=0.0002), SDX 2X (p<0.0061), PYR 2X (p=0.0046), PYR 1/2X (p=0.0313), and SP 2X (0.0279). As stated in the manuscript text, these differences were not consistent between drug concentrations nor even within the same control conditions (S2 Table).

##### Survival

In comparing 14 day survival curves between control un-spiked or solvent-spiked bloodmeals and drug-spiked bloodmeals, amongst multiple comparison analyses of all conditions, there were very limited conditions where survival curves were significantly different: within the AQ/DEAQ feeding group, DEAQ 1/2X condition had worse survival than DEAQ 2X (p=0.0382); within the PPQ feeding group, PPQ 2X and 1X conditions had worse survival than solvent (p=0.0053 and p=0.0413, respectively) (S4 Table).

### Additional statistical results descriptions for *An. coluzzii* mosquito fitness tests

#### Feeding behavior

For mosquito feeding behavior experiments assessing the number of mosquitoes ingesting bloodmeals, beyond analyzing the percent of mosquitoes feeding under any condition, we additionally used chi-squared analysis to compare fed versus unfed mosquitoes for each condition. The chi-squared analyses identified some statistically significant differences comparing mosquitoes fed spiked bloodmeals to those fed an unaltered bloodmeal: mosquitoes within the AQ/DEAQ feeding group fed less with ACN:$H_2O$ 1:9 + FA 0.5% solvent (p<0.0001), IVM 1X (p<0.0001), DEAQ 1X (p=0.023), and DEAQ 1/2X (p<0.0001); mosquitoes within the PPQ feeding group fed less with ACN:$H_2O$ 1:9+FA 0.05% solvent (p<0.0021); and mosquitoes within the SP feeding group fed less for IVM 1X (p<0.0191), SDX 2X (p<0.0001), SDX 1/2X (p=0.0438), PYR 2X (p<0.0001), PYR 1/2X (p=0.0002), and SP 1/2X (p=0.0001), but mosquitoes within the SP feeding group fed more with SP 2X (p=0.0174). This chi-squared analyses also identified some statistically significant differences comparing mosquitoes fed spiked bloodmeals to those fed relevant solvent-spiked bloodmeals: within the AQ/DEAQ feeding group, mosquitoes fed more with DMSO (p=0.0002), AQ 2X (p<0.0001), AQ 1X (p=0.0195), AQ 1/2X (p=0.0004), and DEAQ 2X (p=0.0151); within the PPQ feeding group, mosquitoes fed more with PPQ 1X (p=0.004) and PPQ 1/2X (p=0.0086); and within the SP feeding group mosquitoes fed less with SDX 2X (p<0.0001), PYR 2X (p=0.0106), PYR1/2X (p=0.024), and SP 1/2X (p=0.0104), but within the same SP feeding group fed more with SP 2X (p=0.0002). As stated in the manuscript text, similar to *An. gambiae* results, these differences were not consistent between drug concentrations nor even within the same control conditions (e.g. IVM feeds across experimental groups) (S2 Table).

#### Survival

In comparing 14 day survival curves between control un-spiked or solvent-spiked bloodmeals and drug-spiked bloodmeals, amongst multiple comparison analyses of all conditions, there were very limited conditions where survival curves were significantly different: within the AQ/DEAQ feeding group, blood only control had worse survival compared to the AQ 2X condition (p=0.0279); within the PPQ feeding group, blood only control had worse survival compared to the PPQ 2X (p=0.0069), PPQ 1X (p<0.0001), and solvent (p=0.0002) conditions, and the PPQ 1/2X condition had worse survival than PPQ 1X (p<0.0001) and solvent (p=0.0272) conditions (S4 Table).
