## Supplementary figures and images for "*Anopheles* mosquitoes exposed to long-acting antimalarials via drug-spiked bloodmeal absorb drug but do not suffer fitness costs"

### Supplemental Figure 1

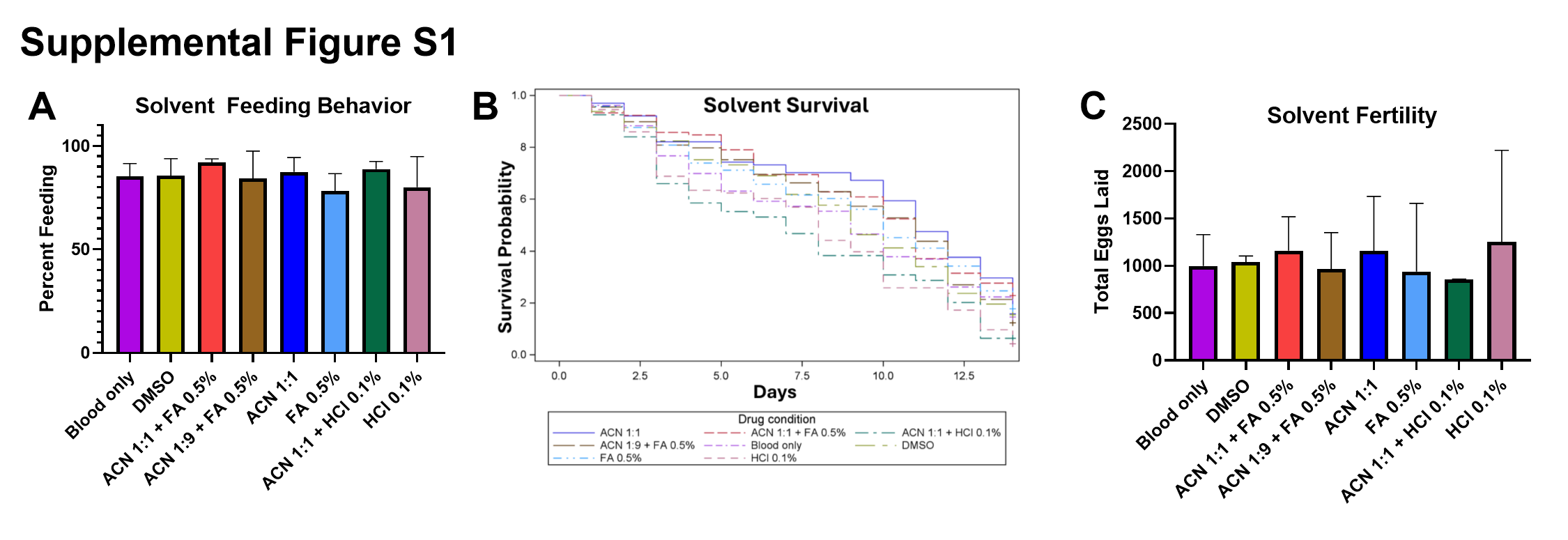

### Supplemental Figure 2

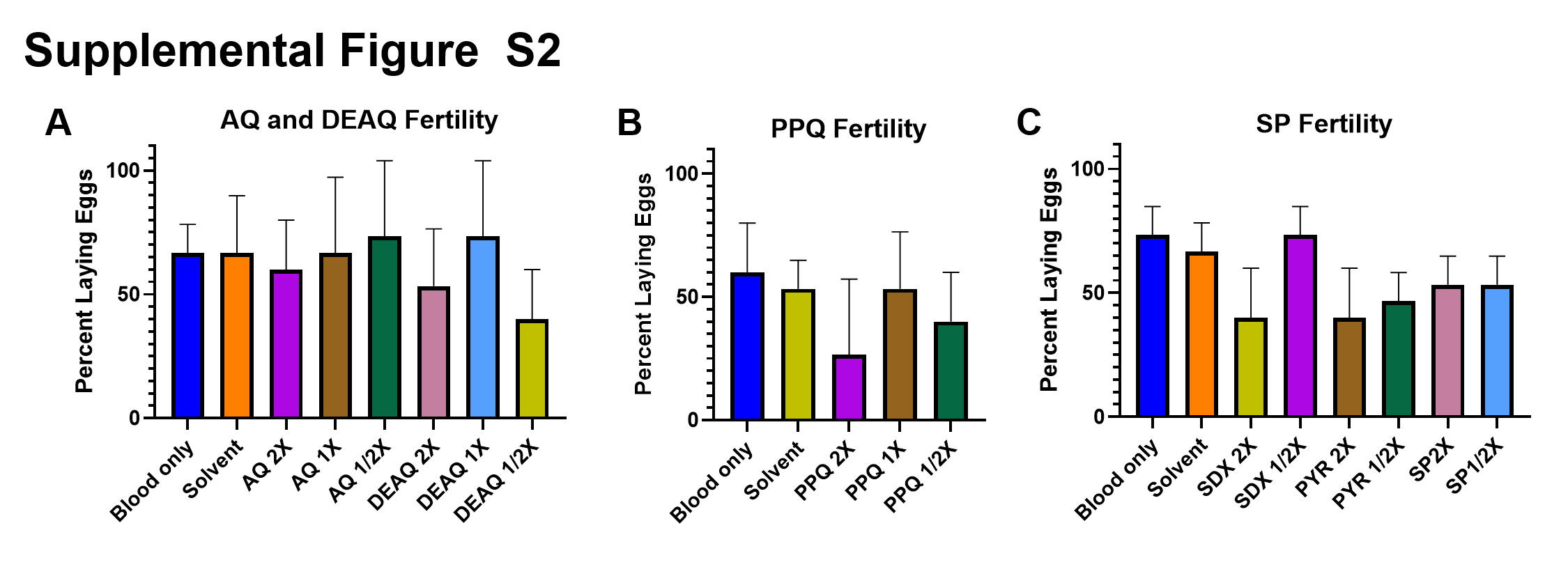

### Supplemental Figure 3

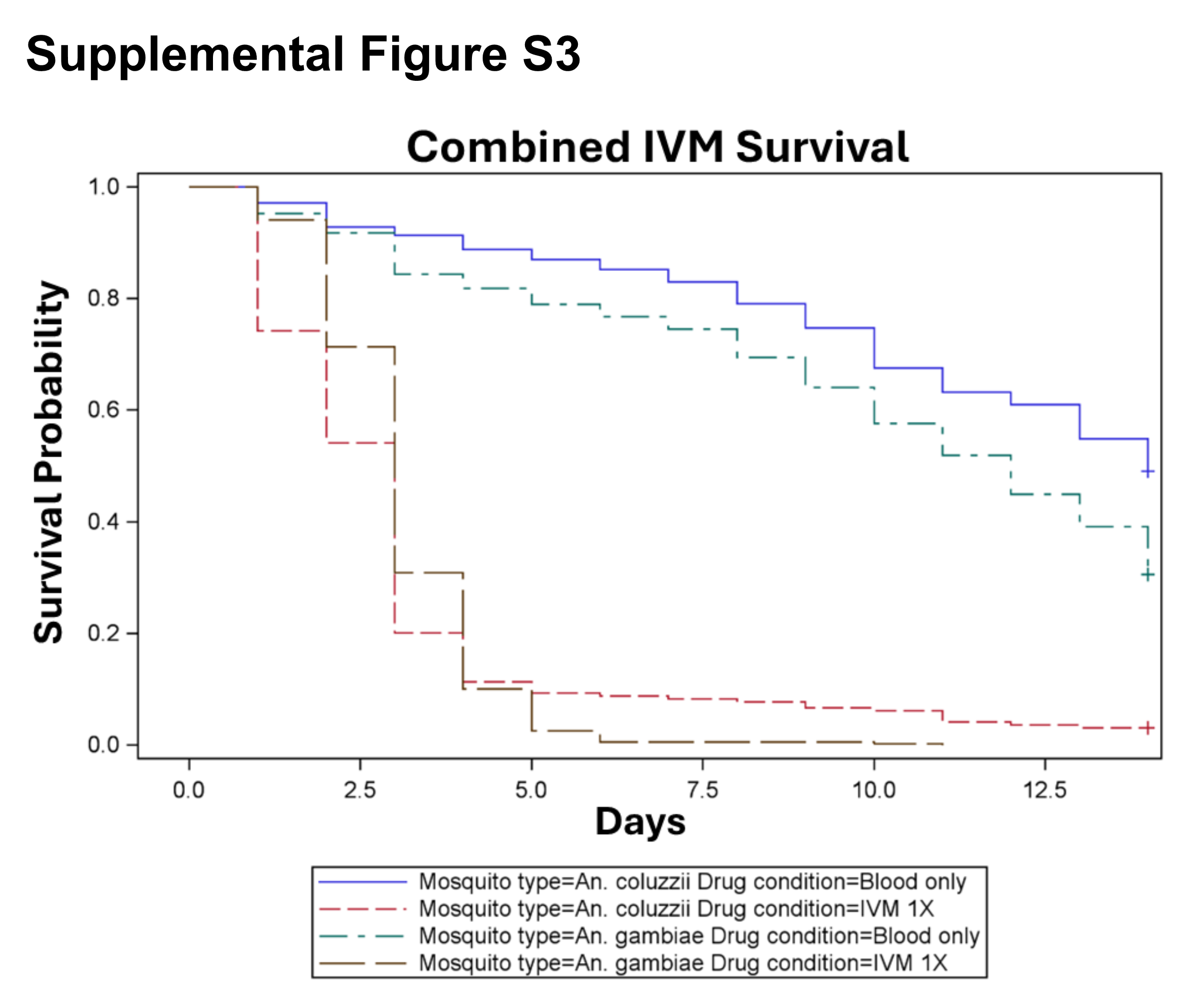

### Supplemental Figure 4

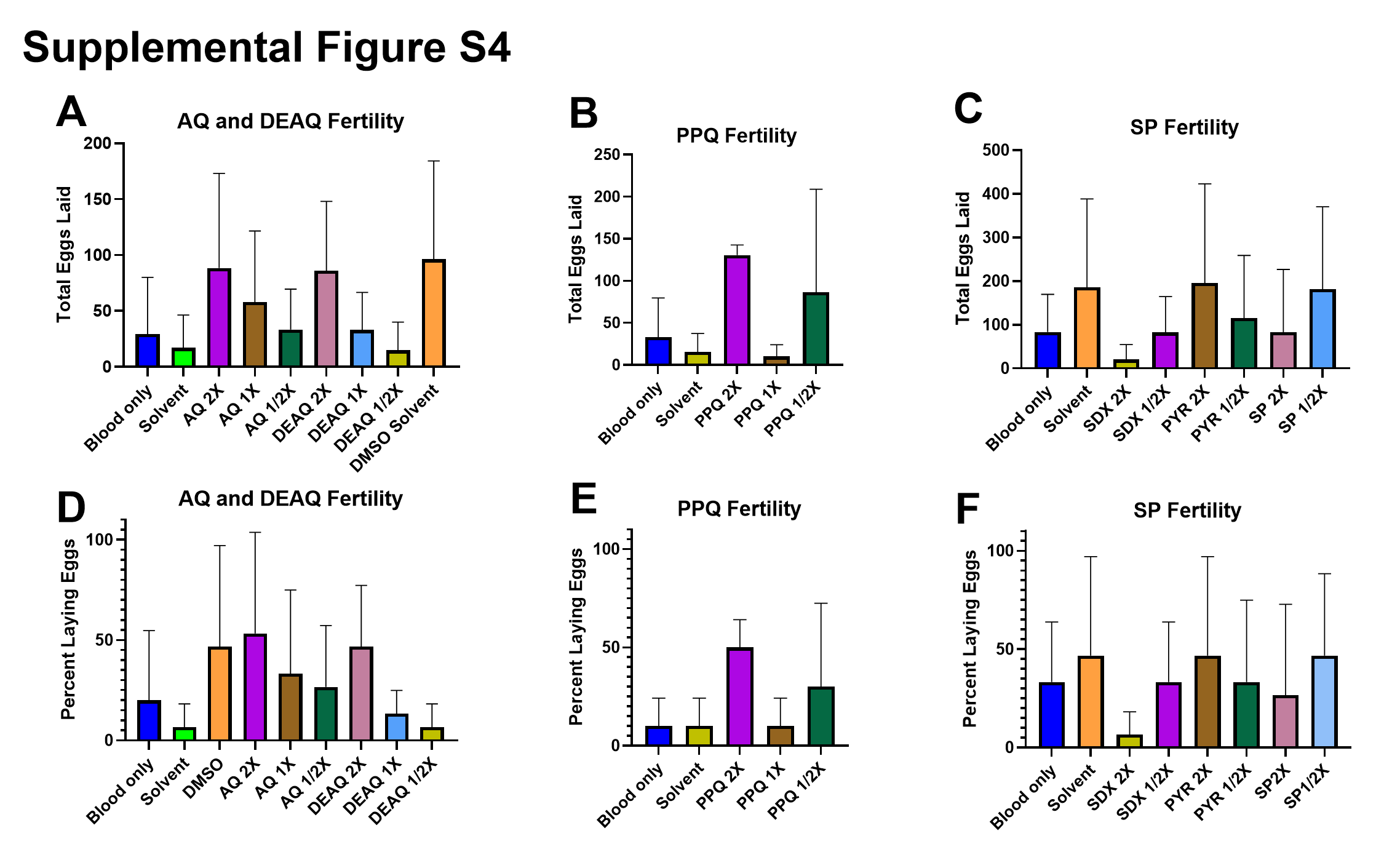
